# Role of TLR4 in persistent *Leptospira interrogans* infection: a comparative *in vivo* study in mice

**DOI:** 10.1101/2020.06.15.153106

**Authors:** Nisha Nair, Mariana Soares Guedes, Adeline Hajjar, Catherine Werts, Maria Gomes-Solecki

**Affiliations:** Department of Microbiology, Immunology and Biochemistry, The University of Tennessee Health Science Center, Memphis, TN, USA; Department of Comparative Medicine, University of Washington, Seattle, Washington, USA; Institut Pasteur, Biology and Genetics of the Bacterial Cell Wall Unit, Paris, France; CNRS, UMR 2001 « Microbiologie intégrative et Moléculaire », Paris, France

**Keywords:** *Leptospira*, leptospirosis, leptospiral-LPS, TLR4, humanized TLR4 mouse, innate immunity, kidney

## Abstract

Toll-Like Receptor (TLR) 4, the LPS receptor, plays a central role in the control of leptospirosis and absence of TLR4 results in lethal infection in mice. Because human TLR4 does not sense the atypical leptospiral-LPS, we hypothesized that TLR4/MD-2 humanized transgenic mice (huTLR4) may be more susceptible to leptospirosis than wild-type mice, and thus may constitute a model of acute human leptospirosis. Therefore, we infected huTLR4 mice, which express human TLR4 but not murine TLR4, with a high but sublethal dose of *L. interrogans* serovar Copenhageni FioCruz (*Leptospira*) in comparison to C57BL/6J wildtype (WT) and, as a control, a congenic strain in which the *tlr4* coding sequences are deleted (muTLR4^Lps-del^). We show that the huTLR4 gene is fully functional in the murine background. We found that dissemination of *Leptospira* in blood, shedding in urine, colonization of the kidney and overall kinetics of leptospirosis progression is equivalent between WT and huTLR4 C57BL/6J mice. Furthermore, inflammation of the kidney appeared to be subdued in huTLR4 compared to WT mice in that we observed less infiltrates of mononuclear lymphocytes, less innate immune markers and no relevant differences in fibrosis markers. Contrary to our hypothesis, huTLR4 mice showed less inflammation and kidney pathology, and are not more susceptible to leptospirosis than WT mice. This study is significant as it indicates that one intact TLR4 gene, be it mouse or human, is necessary to control acute leptospirosis.

**Contribution to the field:** Differences of recognition exist between mouse and human TLR4, in that the anchor of LPS in the outer membrane of *Leptospira* (LipidA) activates murine, but not human TLR4. We hypothesized that if human TLR4 does not sense leptospiral-LPS, then transgenic mice in which murine TLR4 was replaced with human TLR4, should be more susceptible to *Leptospira* dissemination as compared to congenic wild-type mice, which could result in a more robust inflammatory response and pathology in the kidney. However, we found that impaired sensing of leptospiral-LPS in huTLR4 mice did not affect overall infection in comparison to wild-type mice and does not result in increased pathology of the kidney. Our study indicates that rather than leptospiral-LPS sensing, the presence of a fully functional TLR4 receptor is necessary to control acute leptospirosis.

## Introduction

*Leptospira* spp (*Leptospira*) are not considered gram-negative bacteria but they produce lipopolysaccharide (LPS), a potent inflammatory cell wall component that has been shown to be less inflammatory in *Leptospira*. Leptospiral-LPS stimulates mouse and human toll like receptor 2 (TLR2) through abundant associated lipoproteins [1]. Differences of recognition of leptospiral-LPS exist between mouse and human TLR4, in that the anchor of LPS in the outer membrane of *Leptospira* (LipidA) activates murine, but not human TLR4 [1; 2]. This could be due to the lack of phosphate groups in *Leptospira’* LPS lipid A [3]. Escape from human TLR4 may allow for lack of immune recognition, which may favor *Leptospira* infection.

Researchers working on the function of the host TLR4 in leptospirosis used immunocompetent C57BL/6J and its respective congenic TLR4 knockouts. They found that wild-type (WT) mice expressing competent TLR4 receptors in their immune cells are resistant to lethal infection with *L. interrogans* whereas TLR4 knockouts are susceptible, have larger numbers of *Leptospira* in tissues and succumb to infection [4]. Furthermore, we and others [5; 6; 7; 8] developed mouse models of lethal and sublethal leptospirosis using C3H-HeJ mice that have a point mutation in the cytoplasmic domain of the *tlr4* gene [9]. These data strongly suggest that the competence of the TLR4 receptor affects leptospirosis outcomes.

Our hypothesis was that if human TLR4 does not sense leptospiral-LPS [1], then transgenic mice in which murine TLR4 was replaced with human TLR4 (along with its co-receptor MD-2) should be more susceptible to *Leptospira* dissemination as compared to congenic wild-type mice expressing murine TLR4, which could result in a more robust inflammatory response and pathology in the kidney. In this study we compared susceptibility to sublethal leptospirosis in C57BL/6J wild-type (WT) and TLR4 humanized transgenic (huTLR4) mice and evaluated signs of disease progression, as well as pathology and inflammation as we previously described [8; 10]. As a control we used a congenic mutant strain in which the *tlr4* coding sequences are deleted (muTLR4^Lps-del^), as we expected it to develop leptospirosis as shown for other C57BL/6J TLR4^KO^ mice [4].

## Results

To find out if replacing the murine TLR4 with human TLR4 affects the kinetics of leptospirosis progression, we compared sublethal infection in wild-type C57BL/6J (WT) and humanized TLR4/MD2 transgenic mice (huTLR4) and added a congenic murine mutant lacking the TLR4 receptor as control (muTLR4^Lps-del^). We inoculated mice intraperitoneally with a high dose (∼10^8^) of *L. interrogans* Copenhageni FioCruz and analyzed weight loss as a clinical outcome of disease as well as *Leptospira* dissemination and colonization of the kidney (Fig 1 and 2). In contrast to PBS controls, both WT and huTLR4 slowly lost small but significant amounts of weight over two weeks of infection (Fig 1A and 1B). The muTLR4^Lps-del^ lost an abrupt and considerably higher amount of weight after day 6 when colonization of the kidney takes place (Fig 1C). Differences in weight loss between the three infected groups were significant with huTLR4 being the group that lost less weight (Fig 1D).

**Figure 1.**
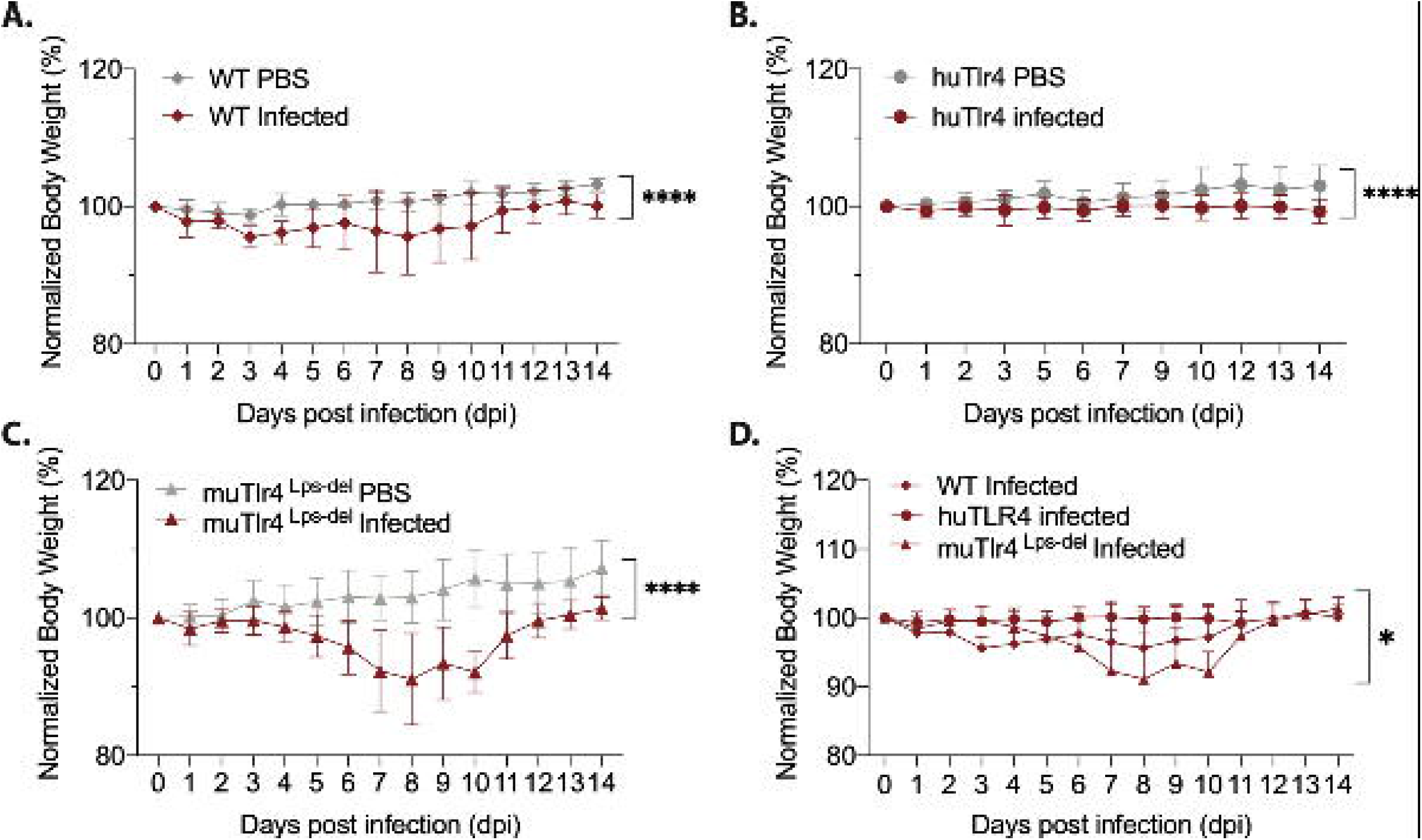
Both WT and huTLR4 mice lose small but significant weight after sublethal infection with *Leptospira interrogans* serovar Copenhageni FioCruz. Weight measurement of C57BL/6 mice carrying wild-type murine TLR4 - WT (A), human TLR4/MD2 - huTLR4 (B) and no murine TLR4 - muTLR4^Lps-del^ - which was used as a control (C). Mice were injected with *Leptospira* (Infected) or with sterile PBS (PBS), n = 7 - 9 per group. Statistics A, B, C by unpaired t test with Welch’s correction, ****p<0.0001 and D, Ordinary One-way ANOVA * p= 0.0168. Data represents two independent experiments.

**Figure 2.**
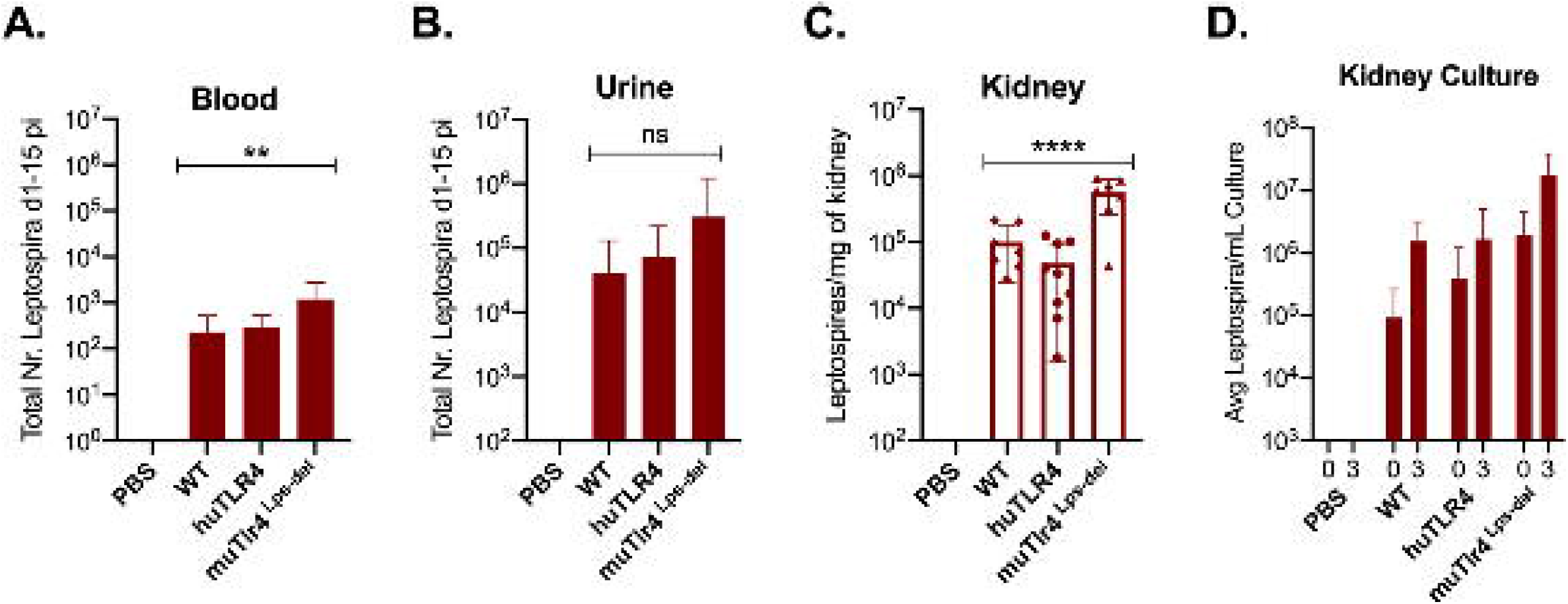
*Leptospira* disseminates in blood and colonizes the kidney of WT and huTLR4 mice. Quantification of total *Leptospira* in blood (A), urine (B), kidney tissue (C), and in kidney culture (D) by qPCR (16S rRNA). Total number (Nr) of *Leptospira* in blood and urine are represented as the sum of *Leptospira* detected between days 1-5 and days 7-14 post-infection, respectively (A and B); kidney is represented as number of *Leptospira* per mg of kidney tissue (C); kidney culture is represented by the average number of *Leptospira* per ml of culture at termination (0) and 3 days later (3) (D). Statistics by one-way ANOVA ** p=0.0074, ****p<0.0001 and ns, non-significant. Data represents two independent experiments.

When we quantified the burden of *Leptospira* in fluids and tissues by qPCR we found low numbers of *Leptospira* in blood in WT and huTLR4 mice (mean ∼230 and ∼300, respectively) and, as expected, a significantly higher number of *Leptospira* in muTLR4 ^Lps-del^ (mean ∼1200) (Fig 2A). Furthermore, much higher numbers of *Leptospira* (∼2-4 Log) were excreted in urine of all infected mice, ∼ 41,000 in WT, ∼74,000 in huTLR4 and ∼305,000 in muTLR4^Lps-del^ (Fig 2B). At termination at 15dpi, tissue PCR showed the same trend with a significantly higher presence of *Leptospira* in the kidney of muTLR4^Lps-del^ (Fig 2C), and culture of the tissue showed that live *Leptospira* colonized the kidney of all groups of infected mice (Fig 2D). These results suggest that *L. interrogans* infection leads to persistent sublethal leptospirosis, renal colonization and stable shedding of *Leptospira* in urine of WT and huTLR4 mice.

To find out if replacing murine TLR4 with human TLR4 affects inflammation of the kidney after infection with *Leptospira*, we analyzed the differences in histopathology and inflammatory markers in tissue collected from infected and uninfected mice two weeks post infection (Fig 3 and 4). Examination of H&E sections of kidney showed that of all three groups of infected mice, huTLR4 had considerably less mononuclear lymphocyte infiltrates and less tubular damage than WT and muTLR4^Lps-del^ (Fig 3A). We purified mRNA from infected and uninfected kidney. Quantification of expression of iNOS and fibroblast activation marker collagen 1 A1 (ColA1) showed an overall lower detection of iNOS in huTLR4 as well as lower ColA1 in WT and huTLR4 than in muTLR4^Lps-del^. Differences between infected groups were statistically significant for ColA1 (Fig 3B).

**Figure 3.**
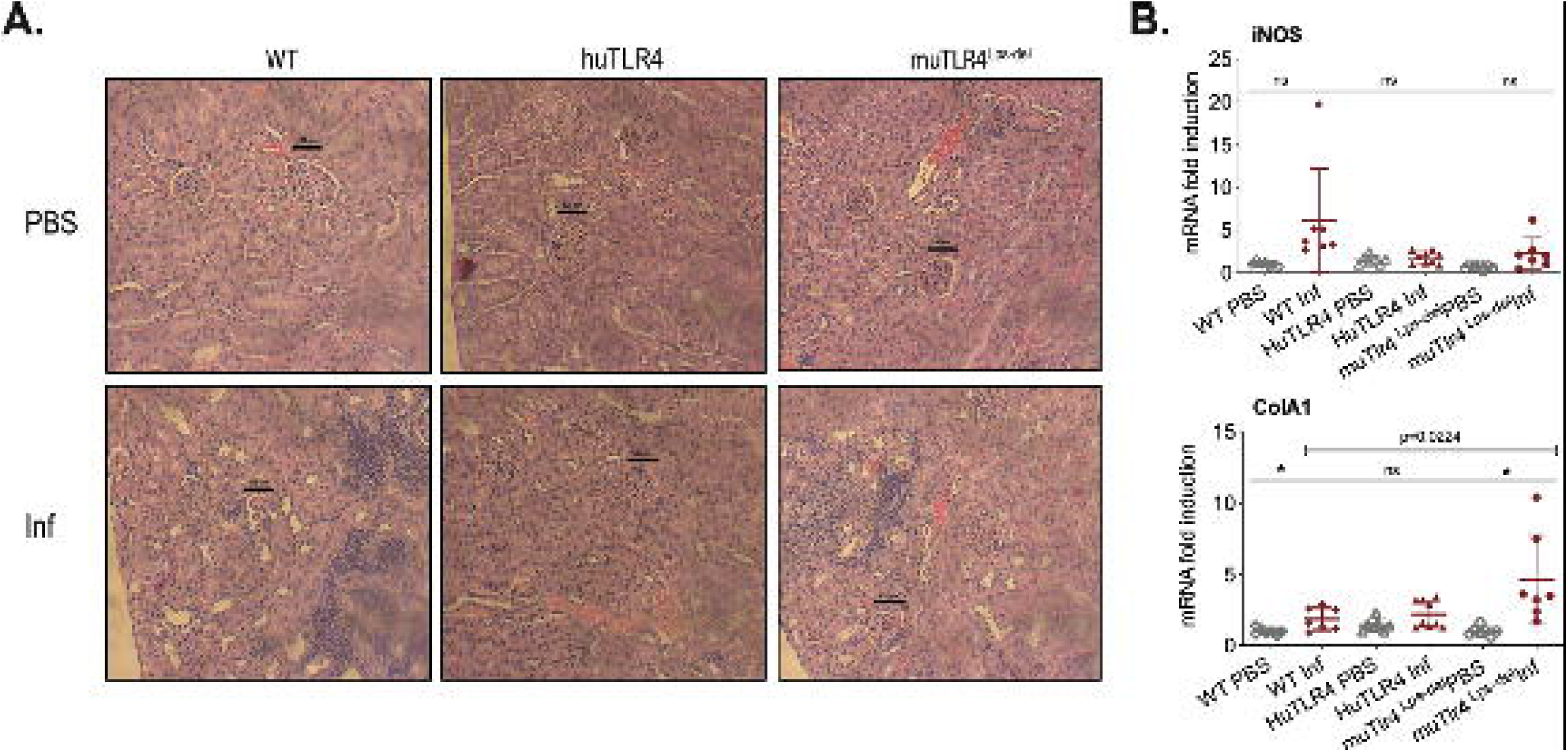
Histopathology and measurements of antimicrobial and collagen markers in the kidney shows less immune cell engagement in huTLR4 mice. A. H&E staining (20X) of kidney sections of *Leptospira* infected (Inf) and uninfected (PBS) C57BL/6J mice carrying wild type murine TLR4 (WT), huTLR4/MD2 (huTLR4) and no murine TLR4 (muTLR4^Lps-del^). Black bar above glomeruli is a 100 micrometer scale and the blue staining highlights infiltration of mononuclear lymphocytes. B. qPCR quantification of antimicrobial transcripts (iNOS) and collagen marker (ColA1). Statistics by unpaired t test with Welch’s correction between infected versus non-infected groups, * p<0.05 and ns, non-significant, and by Ordinary One-Way ANOVA between infected groups for ColA1. H&E data represents one of two experiments and qPCR data represents two independent experiments.

**Figure 4.**
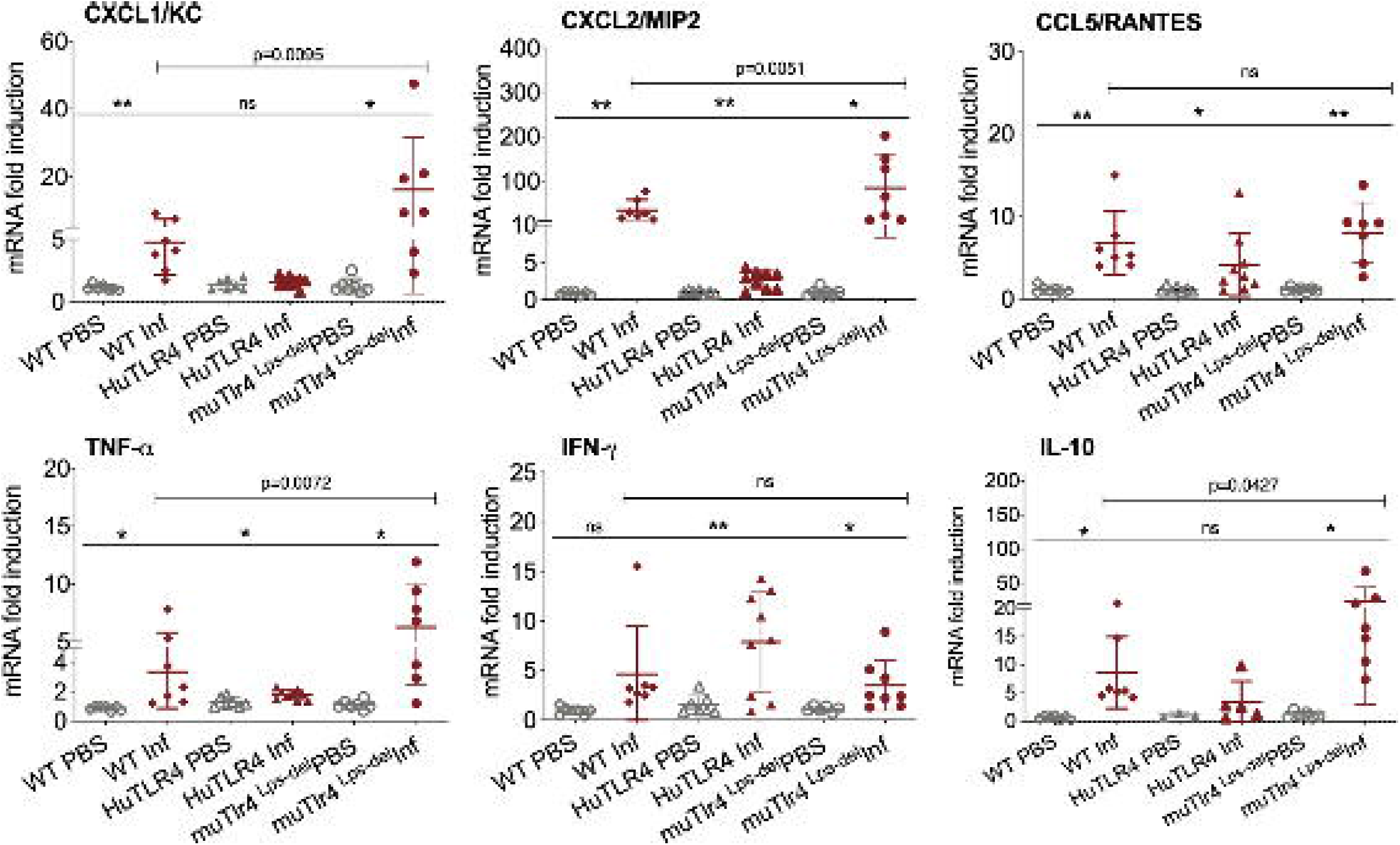
Differences in immune markers in kidney from infected WT and huTLR4 mice suggest less involvement of innate immune processes in huTLR4. qPCR quantification of reverse-transcribed pro-inflammatory innate response chemokines (CxCL1, CxCL2, CCL5) and cytokines (TNF-α, IFNγ, IL-10) in the kidney of wildtype murine TLR4, humanized TLR4 (huTLR4) and muTLR4^Lps-del^ control two weeks post-infection with *Leptospira*. n=7 to 9 mice per group except IL10, n=3 to 7 mice per group. Statistics by unpaired t test with Welch’s correction for infected versus uninfected, * p<0.05, ** p<0.005, and by Ordinary One-Way ANOVA between infected groups, ns, nonsignificant. Data represents two independent experiments.

Given that these results suggest that inflammation may not be triggered in kidney of huTLR4 mice after *Leptospira* infection, we quantified a number of chemokines and cytokines in this tissue (Fig 4). Between infected and uninfected mice, the expression of CxCL1/KC, CxCL2/MIP-2, CCL5/RANTES, TNF-α, IFN-γ and IL-10 were generally significantly increased in the infected kidneys of WT, huTLR4 and muTLR4^Lps-del^ control, with the notable exception of CxCL1/KC chemokine and IL-10 in huTLR4 (Fig 4). Between the three groups of infected mice, differences between CxCL1/KC, CxCL2/MIP2, TNF-α and IL-10 were significant. These results suggest that inflammation in kidneys of huTLR4 mice is reduced compared to WT and muTLR4^Lps-del^ control.

We evaluated systemic splenic cellular immune responses in *Leptospira* infected and noninfected WT, huTLR4 and muTLR4^Lps-del^ mice (Fig 5 and 6). Absolute numbers of splenocytes, CD3+, CD3+CD4+ and CD3+CD8+ T cells were quantified (Fig 5). We found that, although not significant, absolute numbers of splenocytes were increased between infected and uninfected groups. Furthermore, between infected groups, absolute numbers of CD3+, CD3+CD4+ and CD3+CD8+ T cells were reduced in huTLR4 in comparison to WT and muTLR4^Lps-del^ mice.

**Figure 5.**
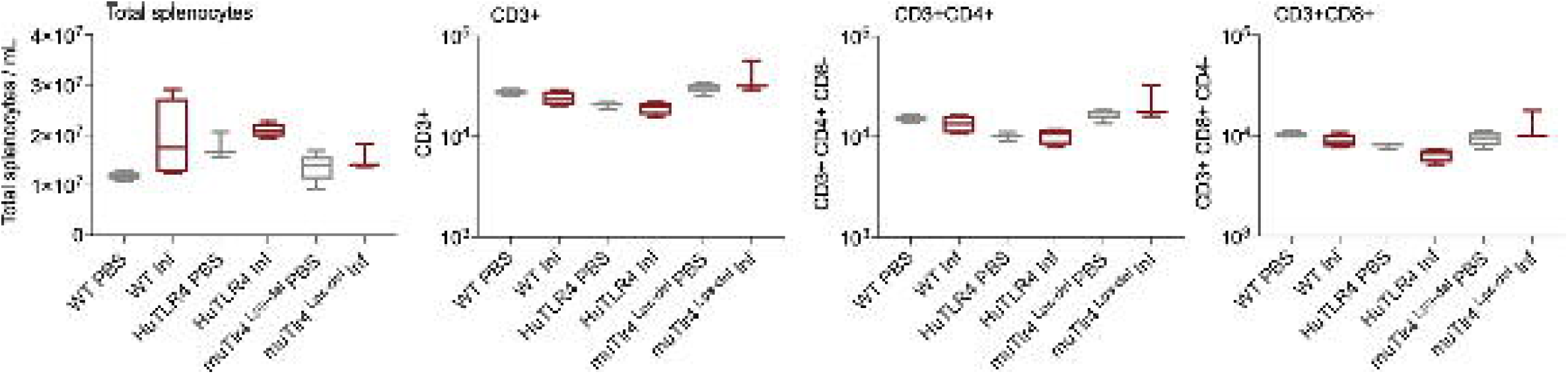
Absolute numbers of total CD3+, CD3+CD4+ and CD3+CD8+ T cells in WT, huTLR4 and muTLR4^Lps-del^ infected with *Leptospira*. Splenocytes were prepared and total cells were counted on a LUNA cell counter. Cells were stained for flow cytometric analysis of T lymphocyte profiles and a total of 100,000 events were captured in a ZE5 cell analyzer (Bio-Rad, Hercules, CA, USA). CD3, CD4, CD8 cell counts were obtained from single cell populations. Data are the mean ±SD of cell counts for 3 or 4 mice per group. Statistics by Unpaired t test with Welch’s correction were not significant between any of the uninfected and infected groups. Data for huTLR4 represents one of two experiments and data for WT/muTLR4^Lps-del^ represents one experiment.

**Figure 6.**
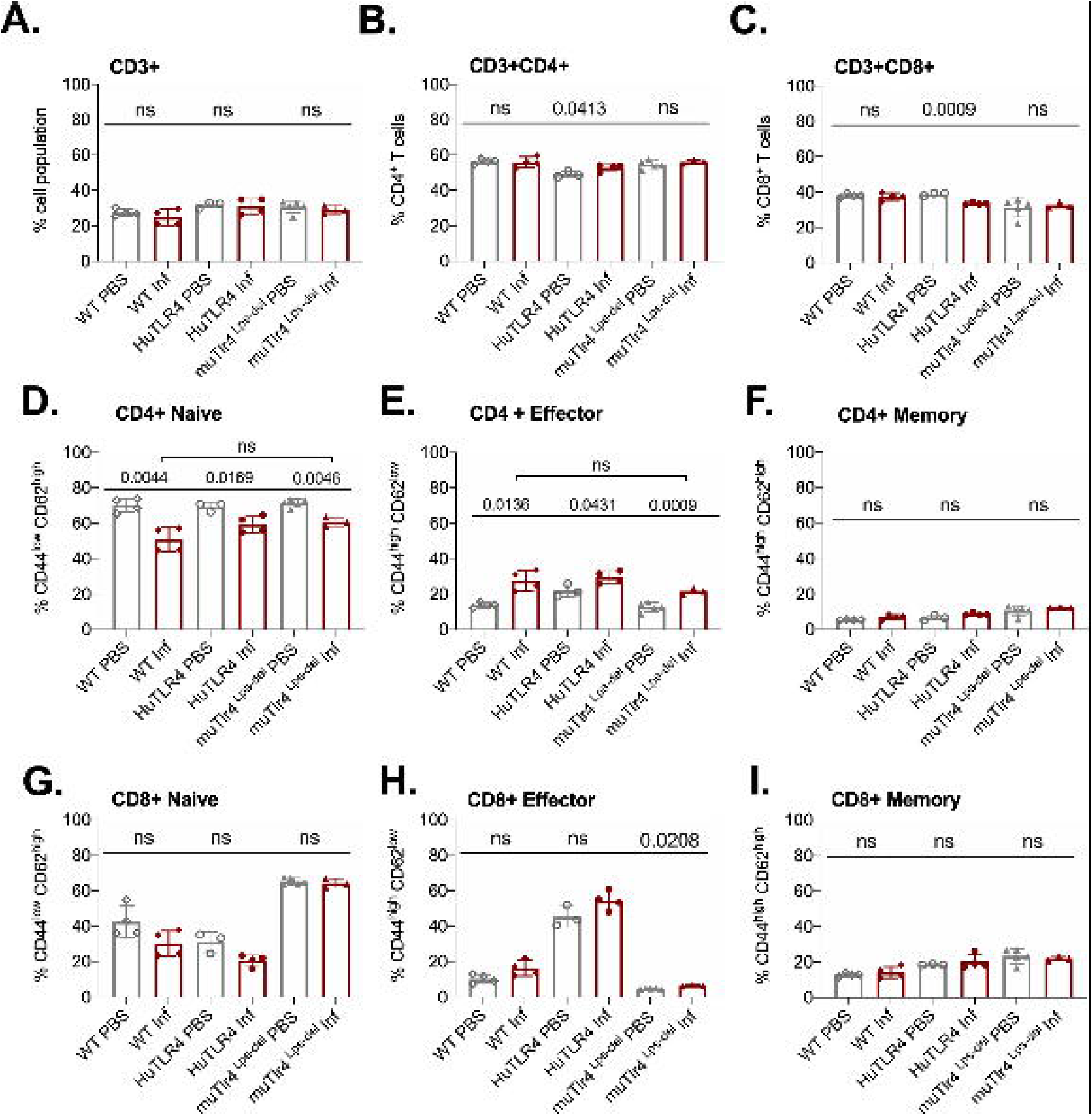
CD4+ effector T cell populations are activated in spleens of WT, huTLR4 and muTLR4^Lps-del^ infected with *Leptospira*. Freshly isolated splenocytes from WT, huTLR4 and muTLR4^Lps-del^ were blocked with anti-mouse CD16/CD32, incubated with the respective surface lineage markers and acquired in a ZE5 cell analyzer (Bio-Rad, Hercules, CA, USA). Data are the mean ±SD percentage of cells for 3 or 4 mice per group. Statistics by Unpaired t test with Welch’s correction between uninfected and infected groups, and by Ordinary One-Way ANOVA between infected groups for CD4+ Naïve and CD4+ Effector; ns, non-significant. Data for huTLR4 represents one of two experiments and data for WT/muTLR4 ^Lps-del^ represents one experiment.

Further flow cytometric analysis of proportions of splenic T cells from WT, huTLR4 and muTLR4^Lps-del^ mice infected with *L. interrogans* (Fig 6) showed no differences in proportions of total CD3+, CD3+CD4+ and CD3+CD8+ T cells between uninfected (PBS) and infected (Inf) mice except for a significantly increased population of CD3+CD4+ and a significantly decreased population of CD3+CD8+ in huTLR4 mice (Fig 6A, 6B, 6C). CD4+ naïve T cell populations decreased between all noninfected and infected groups (Fig 6D) as they differentiated into effector T cells (Fig 6E) but not memory T cells (Fig 6F). Thus, as expected, naïve CD4+ T cells differentiated info effector T cells in infected WT, huTLR4 and muTLR4^Lps-del^ mice. Proportions of CD8+ naïve T cells were slightly lower in infected than in noninfected groups (Fig 6G) as they differentiated into CD8+ effector T cells but not memory T cells. Differences in CD8+ effector T cells were not significant except for muTLR4^Lps-del^ (Fig 6H, 6I).

To get further insight into the role of TLR4 in *Leptospira* induced inflammation, we compared primary splenocytes from WT and huTLR4 mice, as well as mice with a targeted knock-out mutation in TLR4 (muTLR4^KO^) [11]. The cells were stimulated with 10^7^/ml and 10^8^/ml heat-killed *L. interrogans* and gated on the CD11b^+^ macrophage/monocyte population using an *ex vivo* flow-cytometry based assay [12] (Fig 7). We found that 6% and 20% of CD11b+ macrophages from huTLR4 mice stimulated with 10^7^ and 10^8^ *Leptospira*, respectively, produced TNF-α in comparison to ∼ 14% and 28% from WT, and ∼ 3% and 12% from muTLR4^KO^. In addition, macrophages from all mouse genotypes responded to the same extent to CpG and to Pam3CSK4 (TLR9 and TLR2 agonists). WT and huTLR4 macrophages responded to LPS from *E. coli* (EC LPS, TLR4 agonist) in contrast to muTLR4^KO^ macrophages that did not produce TNF-α. It is important to note that stimulation of WT and huTLR4 macrophages with increasing concentrations of the TLR4 agonist (10ng/ml and 100 ng/ml of EC LPS) led to the production of equivalent amounts of TNF-α. This indicates that the huTLR4 receptor is as functional as the WT, in the murine background. Interestingly, previous quantification of relative TLR4 mRNA expression in bone marrow derived macrophages from WT and huTLR4 mice, using the same primers, showed a significantly higher expression of huTLR4 [13]. A lower overall production of TNF-α by huTLR4 macrophages stimulated with 10^7^ and 10^8^ heat-killed *Leptospira* suggests that the systemic inflammatory response of huTLR4 is reduced compared to WT mice.

**Figure 7.**
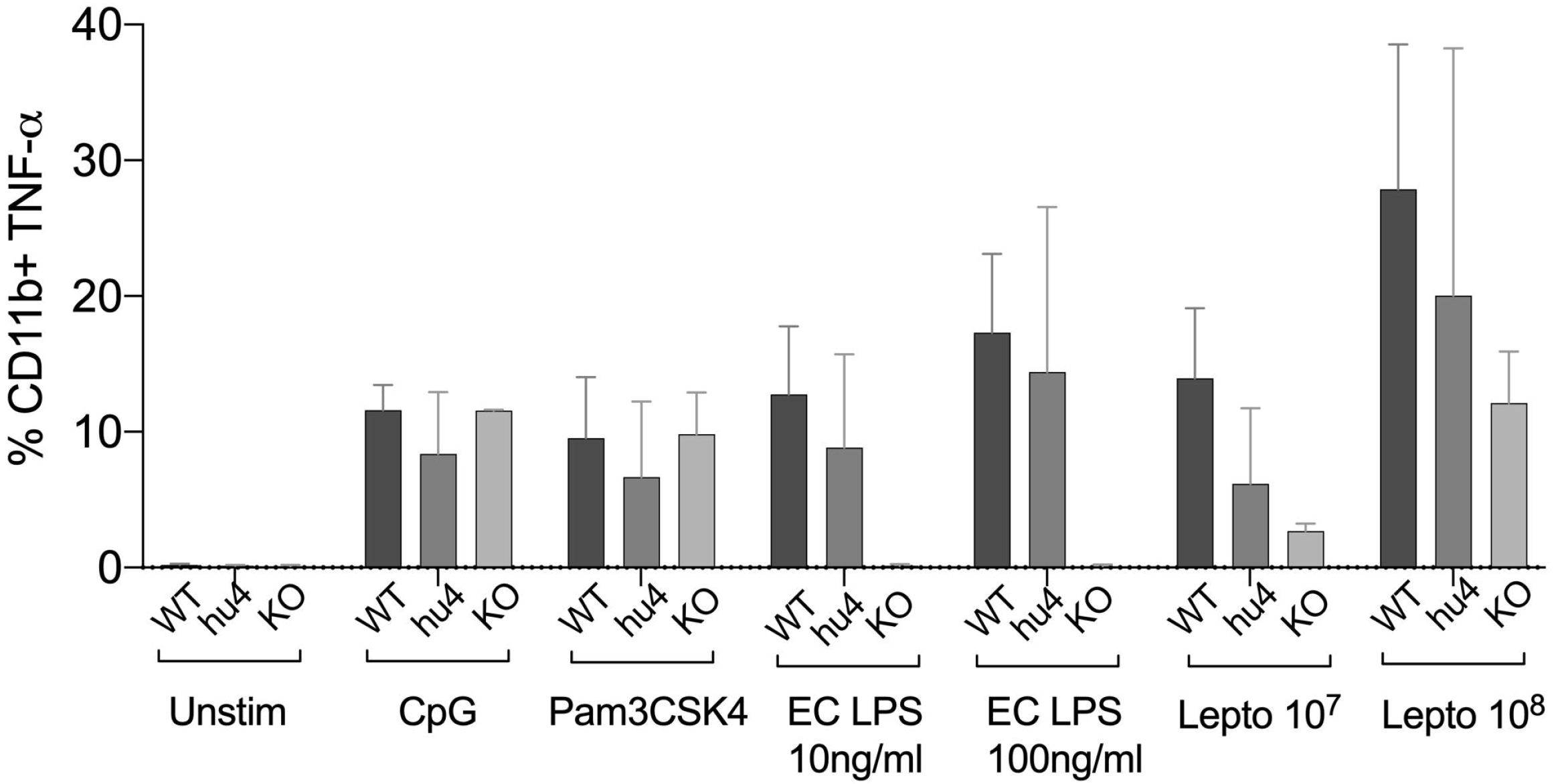
Splenic macrophages of huTLR4 mice produce less TNF-α than WT upon *Leptospira* stimulation. Flow cytometric quantification of % of primary C57BL/6J mouse monocyte/macrophages from WT, huTLR4 and muTLR4^KO^ producing TNF-α (±SD) after stimulation with heat killed *Leptospira* (Lepto, Passage 2) and TLR agonists (CpG ∼ TLR9, Pam3CSK4 ∼TLR2 and *E. coli* (EC) LPS ∼ TLR4). Data represents two experiments.

## Discussion

The major goal of this study was to understand how TLR4 affects *Leptospira* infection *in vivo*. Another goal was to develop an immunocompetent mouse model of persistent or sublethal leptospirosis. The rationale for using humanized TLR4 transgenic mice is that, if human TLR4 does not sense the atypical leptospiral-LPS [14][2][1], these mice should be more permissive of *Leptospira* escape and more susceptible to leptospirosis than wild-type mice expressing murine TLR4 that do sense leptospiral-LPS. We therefore hypothesized that C57BL/6J mice carrying the human TLR4/MD-2 receptor and co-receptor in its immune cells (huTLR4) should be more sensitive to sublethal leptospirosis than wild-type congenic C57BL/6J (WT). However, we found that dissemination of *Leptospira interrogans* serovar Copenhageni FioCruz in blood, shedding in urine, colonization of the kidney and overall kinetics of leptospirosis progression is equivalent between mice carrying murine and human TLR4. Furthermore, the inflammatory response to infection with *L. interrogans* seemed to be subdued in huTLR4 compared to WT mice in that we observed less infiltrates of mononuclear lymphocytes, less innate immune markers and no relevant differences in fibrosis markers in kidney as well as less production of TNFα by murine splenic macrophages harboring a fully functional huTLR4 receptor. These results are consistent with the lack of leptospiral-LPS stimulation of human TLR4 [1].

Decreased inflammation in huTLR4 mice could have led to increased numbers of *Leptospira* in kidney. Surprisingly, this was not observed. We analyzed the kinetics of leptospirosis infection and colonization of the kidney between infected and uninfected humanized TLR4 mice as established previously in our laboratory [8][10][15] (Fig 1 and 2). Compared to WT and to muTLR4^Lps-del^, infected huTLR4 mice showed a similar pattern of low and slow loss of weight as the WT which was considerably different than the abrupt loss of weight shown by muTLR4^Lps-del^ at days ∼6-7 post-infection when colonization of the kidney takes place, as we previously observed in C3H-HeJ mice [8][10][16][15]. The latter carries a point mutation in the toll like receptor 4 (*tlr4*) cytoplasmic domain that results in a lack of signaling despite presence of the receptor and binding of bacterial LPS [9]. Furthermore, dissemination in blood and shedding of *Leptospira* in urine of WT and huTLR4 mice were equivalent as was the amount of live *Leptospira* detected in kidney. This again contrasted with the amount of *Leptospira* detected in blood, urine and kidney of muTLR4^Lps-del^ mice that were 1Log higher than the WT and huTLR4 counterparts. An even higher load of *Leptospira* was observed in blood and urine of infected C3H-HeJ mice [8][15]. Furthermore, wild-type C57BL/6J mice were found to have 2Log lower numbers of *Leptospira* in urine than TLR4^KO^ mice [4][17]. Interestingly, the results of *Leptospira* infection obtained in C3H-HeJ (containing a point mutation in the TLR4 receptor) and in C57BL/6J TLR4^KO^ (do not express the TLR4 receptor) are consistent with the results obtained here with C57BL/6.10J TLR4^Lps-del^ which contain a spontaneous mutation that completely removes the *tlr4* gene and thus results in the absence of the TLR4 receptor. These results confirm that mutations resulting in a loss of response to leptospiral-LPS, either resulting from a point mutation in *tlr4* in C3H-HeJ or from a complete deletion of the gene in C57BL6/J mice, increase susceptibility to *Leptospira* infection regardless of genetic background. However, this was not replicated by substitution of wildtype murine TLR4 with human TLR4, that is known to lack leptospiral-LPS sensing.

Analysis of markers of inflammation and other immune cell engagement (Fig 3 and 4) in kidney showed that huTLR4 mice exhibit less markers of innate immune cell activity (CxCL1/KC, CxCL2/MIP-2 and TNF-α) and less anti-inflammatory cytokine IL-10 than WT mice. Differences in antimicrobial iNOS and fibroblast activation marker collagen 1 A1 (ColA1) which are indicative of interstitial collagen deposition (fibrosis) were not significant between huTLR4 and WT mice. This may be further corroborated by the lower number of mononuclear lymphocyte infiltrates observed in H&E stained slides of the kidney. These data suggest that decreased inflammatory responses in the huTLR4 mouse kidney is dependent on TLR4. If human TLR4 does not sense the atypical leptospiral-LPS [14][2][1], huTLR4 mice should have been more permissive of *Leptospira* escape and more susceptible to leptospirosis than wild-type. However, the opposite was observed. The data could suggest that the huTLR4 transgene may not function to their fullest capacity in a mouse background as it does in humans. However, our data from splenic murine macrophages stimulated with *Leptospira* (Fig 7) clearly shows that the huTLR4 receptor is fully functional in mouse macrophages as they produced as much TNF-α after stimulation with increasing doses of the *E. coli* LPS TLR4 agonist as did WT mice. Although stimulation of TLR9 and TLR2 led to equivalent production of TNF-α by WT, huTLR4 and muTLR4^Lps-del^ mouse macrophages, recognition of heat killed *Leptospira* was decreased in huTLR4 when compared to WT, which is consistent with the lack of sensing of leptospiral-LPS by human TLR4 [1].

When we analyzed proportions of splenic T cells two weeks post infection with *L. interrogans* (Fig 5 and 6) we found increased populations of CD4+ effector T cells in infected versus uninfected mice in WT, huTLR4 and muTLR4^Lps-del^ which was consistent with our previous observations in C3H-HeJ mice [8][10]. Furthermore, the differences observed in CD3+CD8+ T cells suggest that cytotoxic CD8+ T cells could be engaged in mounting an immune response to *Leptospira*. Interestingly, in cattle vaccinated with a *Leptospira* bacterin and subsequently challenged with *L. borgpetersenii* serovar Hardjo, CD8+ were the only T cells that proliferated 6 weeks post infection [18]. Thus, independently of genetic background, effector CD4+ T cells promote the T cell response to *L. interrogans* infection and, quite possibly, CD8+ T cells may be involved.

Regarding differences in infection in immunocompetent (C57BL/6J WT) versus immunocompromised mice (C3H-HeJ, and C57BL/6.10J TLR4^Lps-del^ and C57BL/6J TLR4^KO^), *Leptospira* persistence in WT mice, shown in ours and other studies [17] fulfils their role as host reservoirs. In the current study we show that *Leptospira* dissemination, although lower in WT C57BL/6J than in C3H-HeJ [8], can be quantified and the immune marker profiles of infection are similar between the two strains. The differences in order of magnitude observed between the two strains simply allow for a more robust quantification of dissemination and inflammation in C3H-HeJ [8; 15]. It is however important to note that *Leptospira* disseminated and caused pathology in wild-type C57BL/6J which carry a fully functional murine TLR4. This could be explained by the recent finding that leptospiral-LPS also subverts murine TLR4 recognition [19]. Indeed, leptospiral-LPS recognition by murine TLR4 is only partial, since only the MyD88 pathway is engaged, not the endosomal TRIF pathway. This avoidance is due to LPS-associated lipoproteins that interfere with CD14 binding of leptospiral-LPS, which block the internalization of TLR4. As a consequence of TRIF avoidance, secretion of type I interferons and nitric oxide production upon *Leptospira* infection are decreased [19]. This somewhat altered TLR4 response could explain the dissemination and kidney colonization after *Leptospira* infection in mice and it can provide a clue about the lack of profound differences between huTLR4 and WT mice after infection. In addition, *Leptospira* evade different PRRs, such as TLR5 and NOD proteins, which could contribute to the poor efficiency of macrophages and phagocytes facing *Leptospira* [20]. In this context, the overall role of TLR4 sensing *Leptospira* could be minimal although its presence would still be important. For example, TLR4 expressed by B cells was shown to be key in early IgM production in mice, which is partially able to counteract *Leptospira* infection [4]. We may speculate that early IgM production, activated three days post infection, may be linked to a role of TLR4, as it was recently shown that B1 cell activation and tumor-reactive IgM production were defective in TLR4^KO^ mice [21].

In conclusion, we expected huTLR4 mice to have been more permissive of *Leptospira* escape and more susceptible to leptospirosis than congenic wild-type mice. Our hypothesis did not prove to be true. Our data suggests that less recognition of leptospiral-LPS by huTLR4 did not affect overall infection and does not result in increased pathology of the kidney. These data indicate that an intact TLR4 gene, be it mouse or human, is necessary to control pathology caused by *Leptospira*. Unexpectedly, this could help explain why ∼90% of people infected with pathogenic *Leptospira* [22] are fairly tolerant of this pathogen and do not develop acute Weil’s disease.

## Materials and Methods

### Bacterial strains

We used *Leptospira interrogans* serovar Copenhageni strain Fiocruz L1-130 (*Leptospira*), culture passage 2 after hamster infection, originally isolated from a patient in Brazil. *Leptospira* was cultured as previously described [16] and enumerated by dark-field microscopy (Zeiss USA, Hawthorne, NY) that was confirmed by qPCR (StepOne Plus, Life Technologies, Grand Island, NY).

### Animals

TLR4/MD-2 humanized transgenic mice (huTLR4) were developed by Dr Hajjar on a C57BL/6J TLR4/MD-2 double knock-out background [11; 12; 23]. 6-week old huTLR4 male mice were transferred from Dr Hajjar laboratory at the University of Washington to the Laboratory Animal Care Unit of the University of Tennessee Health Science Center (LACU-UTHSC) and quarantined for 3 weeks. C57BL/6J mice with a targeted mutation in TLR4 to produce a null (muTLR4^ko^) [11], were used to derive primary macrophages to stimulate with heat-killed *Leptospira* and other TLR agonists to verify the receptor function. Male 9-week old wild-type C57BL/6J (WT) mice and a congenic C57BL/6.10J mutant strain B6.B10ScN-Tlr4lps-del/JthJ, stock #007227 [24; 25], (muTLR4^Lps-del^) were purchased from The Jackson Laboratories (Bar Harbor, ME) and acclimatized for one week at the pathogen-free environment in the LACU-UTHSC. The mutant strain (muTLR4^Lps-del^) contains a spontaneous mutation that completely removes the *tlr4* gene which results in absence of the TLR4 protein. Infections were done when mice reached 10 weeks of age.

### Ethics statement

*This study was carried out in accordance with the Guide for the Care and Use of Laboratory Animals of the NIH*. The protocols were approved by the University of Tennessee Health Science Center (UTHSC) Institutional Animal Care and Use Committee, Animal Care Protocol Application, Permits Number 16-070 and 19-062.

### Protocols

Comprehensive protocols for infection of mice with pathogenic *Leptospira*, for monitoring clinical and molecular scores of disease and for evaluation of the host immune response were published recently [26].

### Infection of mice

Intraperitoneal infection was done as described previously using a dose of ∼10^8^ virulent *Leptospira* in sterile PBS. Bacteria were counted in a Petroff-Hausser chamber under a dark field microscope and confirmed by qPCR. Body weights were monitored daily. Urine was also collected on a daily basis for 15 days p.i. by gently massaging the bladder area and the urine was collected into sterile aluminum foil. Blood (up to 20 μl) was collected every other day by tail nick for 15 days. At termination, kidneys were collected for qPCR.

### RT-PCR and q-PCR

DNA was extracted per manufacturers’ instructions from urine, blood, and kidney using a NucleoSpin tissue kit (Clontech). Quantification of *Leptospira* 16s rRNA was done using TAMRA probe and primers from Eurofins (Huntsville, AL) by real-time PCR (qPCR) (StepOne Plus). RNeasy mini kit (Qiagen) was used to extract total RNA followed by reverse transcription using a high-capacity cDNA reverse transcription kit (Applied Biosystems). Real-time PCR on the cDNA was performed as described [8]. For RT-PCR, we used TAMRA probes specific for inducible nitric oxide synthase (iNOS), Collagen A1 (ColA1), keratinocyte-derived chemokine (KC, CxCL1), macrophage inflammatory protein 2 (MIP-2, CxCL2), RANTES (CCL5), tumor necrosis factor alpha (TNF-α), interferon gamma (IFN-γ) and IL-10. β-actin was used as control for the comparative CT method [10].

### Histopathology

The kidney tissues were fixed in 10% neutral buffered formalin and submitted to the UTHSC Research Histology Core Facility for histology processing. Paraffin embedded tissues were sectioned and stained with hematoxylin and eosin (H&E). The slides were analyzed for nephritis and imaged under an Axio Zeiss Imager A1 light microscope with Axiocam 103 and Zen 2 lite software.

### Profiling immune cell populations (T and B cells) by Flow Cytometry

Cell preparation: Spleens were harvested from mice, placed in RPMI 1640 (Mediatech, Manassas, VA), and diced with frozen slides in Hanks balanced salt solution medium (Cellgro, Manassas, VA) to produce cell suspensions. Red blood cell lysis was performed using 2 ml/spleen of 1X ACK lysing buffer (Life Technologies). Cells were then washed in RPMI 1640 containing 10% heat-inactivated fetal bovine serum (Atlanta Biotech) and 0.09% sodium azide (Sigma-Aldrich, St. Louis, MO), followed by passage through 70-μm-pore-size and 40-μm-pore-size nylon filters (BD Falcon, Bedford, MA), and the cells were counted using a Luna-FL automated cell counter (Logos Biosystems, Gyunggi-Do, South Korea).

Flow Cytometric Cell Staining: Cell viability was analyzed by mixing 18 μl of splenocytes with 2 μl of acridine orange-propidium iodide staining solution in a Luna-FL automated cell counter, and 3×10^6^ cells per tube were stained as follows: cells were incubated in 0.5 μg Fc block (BD Biosciences) for 15 min at 4°C in staining buffer (Ca^2+^ - and Mg^2+^ - free PBS containing 3% heat-inactivated fetal bovine serum, 0.09% sodium azide [Sigma-Aldrich, St. Louis, MO], 5mM EDTA), washed twice, and incubated with the appropriate marker for surface staining in the dark for 30 min at 4°C. T lymphocyte staining was performed using CD3 clone 17A2 conjugated with fluorescein isothiocyanate (FITC) (1:200; BD Biosciences), CD4 clone RM4-5 conjugated with phycoerythrin (PE) (1:150; BD Biosciences), CD8 clone 53-6.7 conjugated with allophycocyanin (APC)-Cy7 (1:150; BD Biosciences), CD62L clone MEL-14 conjugated with PE-Cy7 (1:150; BD Biosciences), CD44 clone IM7 conjugated with APC (1:150; BD Biosciences). Flow cytometric staining and analysis was done as described [8] and acquired on ZE5 cell analyzer (Bio-Rad, Hercules, CA, USA). Analysis was performed using Flow Jo software.

### Splenic macrophages

Heat-killed *Leptospira* were prepared as follows: pellets from a culture, which were washed with 1 ml PBS, were harvested by centrifugation at 12,000g and heat killed by exposure to heat for 10 min at 95°C. The supernatant was discarded, and the extract resuspended in endotoxin free PBS. Lack of motility was confirmed under a dark field microscope. This process was repeated to ensure no motile *Leptospira* were visualized.

Single-cell splenocyte suspensions were stimulated with 10 μg/ml CpG 1826 or Pam3CSK4, 10 ng/ml and 100 ng/ml *E. coli* O111:B4 LPS, or 10^7^/ml and 10^8^/ml heat-killed P2 *Leptospira* in the presence of 10 μg/ml brefeldin A for ∼ 4hr as described [12]. Cells were then stained and analyzed using flow cytometry as described [12]. CpG, *E. coli* (EC) LPS and Pam3cys were used as TLR9, TLR4 and TLR2 agonists, respectively.

### Statistics

Two-tailed unpaired parametric t-test with Welch’s correction was used to analyze differences between infected and uninfected groups. Differences between infected mice was analyzed by Ordinary One-Way ANOVA. For immune markers and T cell populations, Ordinary One-Way ANOVA was done if at least two of the mouse strains showed significant differences between infected and uninfected mice to ensure the effect seen was due to infection. Statistical analysis was done using GraphPad Prism software, α = 0.05.

## Acknowledgements

We thank the Flow Cytometry and Cell Sorting Core at UTHSC. This work was supported by Public Health Service grants R44AI096551, R43AI136551, R21AI142129 (M.G.S.) and from the National Institutes of Health and supported by an Institut Pasteur grant PTR66-2017 (CW).

## References

[1] M.A. Nahori, E. Fournie-Amazouz, N.S. Que-Gewirth, V. Balloy, M. Chignard, C.R. Raetz, I. Saint Girons, and C. Werts, Differential TLR recognition of leptospiral lipid A and lipopolysaccharide in murine and human cells. J Immunol 175 (2005) 6022–31.

[2] N.L. Que-Gewirth, A.A. Ribeiro, S.R. Kalb, R.J. Cotter, D.M. Bulach, B. Adler, I.S. Girons, C. Werts, and C.R. Raetz, A methylated phosphate group and four amide-linked acyl chains in leptospira interrogans lipid A. The membrane anchor of an unusual lipopolysaccharide that activates TLR2. J Biol Chem 279 (2004) 25420–9.

[3] T. Scior, C. Alexander, and U. Zaehringer, Reviewing and identifying amino acids of human, murine, canine and equine TLR4 / MD-2 receptor complexes conferring endotoxic innate immunity activation by LPS/lipid A, or antagonistic effects by Eritoran, in contrast to species-dependent modulation by lipid IVa. Comput Struct Biotechnol J 5 (2013) e201302012.

[4] C. Chassin, M. Picardeau, J.M. Goujon, P. Bourhy, N. Quellard, S. Darche, E. Badell, M.F. d’Andon, N. Winter, S. Lacroix-Lamande, D. Buzoni-Gatel, A. Vandewalle, and C. Werts, TLR4- and TLR2-mediated B cell responses control the clearance of the bacterial pathogen, Leptospira interrogans. J Immunol 183 (2009) 2669–77.

[5] M.M. Pereira, J. Andrade, R.S. Marchevsky, and R. Ribeiro dos Santos, Morphological characterization of lung and kidney lesions in C3H/HeJ mice infected with Leptospira interrogans serovar icterohaemorrhagiae: defect of CD4+ and CD8+ T-cells are prognosticators of the disease progression. Exp Toxicol Pathol 50 (1998) 191–8.

[6] J.E. Nally, M.C. Fishbein, D.R. Blanco, and M.A. Lovett, Lethal infection of C3H/HeJ and C3H/SCID mice with an isolate of Leptospira interrogans serovar copenhageni. Infect Immun 73 (2005) 7014–7.

[7] S. Viriyakosol, M.A. Matthias, M.A. Swancutt, T.N. Kirkland, and J.M. Vinetz, Toll-like receptor 4 protects against lethal Leptospira interrogans serovar icterohaemorrhagiae infection and contributes to in vivo control of leptospiral burden. Infect Immun 74 (2006) 887–95.

[8] L. Richer, H.H. Potula, R. Melo, A. Vieira, and M. Gomes-Solecki, Mouse model for sublethal Leptospira interrogans infection. Infect Immun 83 (2015) 4693–700.

[9] S.N. Vogel, D. Johnson, P.Y. Perera, A. Medvedev, L. Lariviere, S.T. Qureshi, and D. Malo, Cutting edge: functional characterization of the effect of the C3H/HeJ defect in mice that lack an Lpsn gene: in vivo evidence for a dominant negative mutation. J Immunol 162 (1999) 5666–70.

[10] H.H. Potula, L. Richer, C. Werts, and M. Gomes-Solecki, Pre-treatment with Lactobacillus plantarum prevents severe pathogenesis in mice infected with Leptospira interrogans and may be associated with recruitment of myeloid cells. PLoS Negl Trop Dis 11 (2017) e0005870.

[11] K. Hoshino, O. Takeuchi, T. Kawai, H. Sanjo, T. Ogawa, Y. Takeda, K. Takeda, and S. Akira, utting edge: Toll-like receptor 4 (TLR4)-deficient mice are hyporesponsive to lipopolysaccharide: evidence for TLR4 as the Lps gene product. J Immunol 162 (1999) 3749–52.

[12] A.M. Hajjar, R.K. Ernst, E.S. Fortuno, 3rd, A.S. Brasfield, C.S. Yam, L.A. Newlon, T.R. Kollmann, S.I. Miller, and C.B. Wilson, Humanized TLR4/MD-2 mice reveal LPS recognition differentially impacts susceptibility to Yersinia pestis and Salmonella enterica. PLoS Pathog 8 (2012) e1002963.

[13] A.M. Hajjar, R.K. Ernst, J. Yi, C.S. Yam, and S.I. Miller, Expression level of human TLR4 rather than sequence is the key determinant of LPS responsiveness. PLoS One 12 (2017) e0186308.

[14] C. Werts, R.I. Tapping, J.C. Mathison, T.H. Chuang, V. Kravchenko, I. Saint Girons, D.A. Haake, P.J. Godowski, F. Hayashi, A. Ozinsky, D.M. Underhill, C.J. Kirschning, H. Wagner, A. Aderem, P.S. Tobias, and R.J. Ulevitch, Leptospiral lipopolysaccharide activates cells through a TLR2-dependent mechanism. Nat Immunol 2 (2001) 346–52.

[15] N. Nair, M.S. Guedes, C. Werts, and M. Gomes-Solecki, The route of infection with Leptospira interrogans serovar Copenhageni affects the kinetics of bacterial dissemination and kidney colonization. PLoS Negl Trop Dis 14 (2020) e0007950.

[16] J.P. Sullivan, N. Nair, H.H. Potula, and M. Gomes-Solecki, Eye-Drop Inoculation Leads to Sublethal Leptospirosis in Mice. Infect Immun (2017).

[17] M. Fanton d’Andon, N. Quellard, B. Fernandez, G. Ratet, S. Lacroix-Lamande, A. Vandewalle, I.G. Boneca, J.M. Goujon, and C. Werts, Leptospira Interrogans induces fibrosis in the mouse kidney through Inos-dependent, TLR- and NLR-independent signaling pathways. PLoS Negl Trop Dis 8 (2014) e2664.

[18] R.L. Zuerner, D.P. Alt, M.V. Palmer, T.C. Thacker, and S.C. Olsen, A Leptospira borgpetersenii serovar Hardjo vaccine induces a Th1 response, activates NK cells, and reduces renal colonization. Clin Vaccine Immunol 18 (2011) 684–91.

[19] D. Bonhomme, I. Santecchia, F. Vernel-Pauillac, M. Caroff, P. Germon, G. Murray, B. Adler, I.G. Boneca, and C. Werts, Leptospiral LPS escapes mouse TLR4 internalization and TRIFassociated antimicrobial responses through O antigen and associated lipoproteins. PLoS Pathog 16 (2020) e1008639.

[20] I. Santecchia, M.F. Ferrer, M.L. Vieira, R.M. Gomez, and C. Werts, Phagocyte Escape of Leptospira: The Role of TLRs and NLRs. Front Immunol 11 (2020) 571816.

[21] A.M. Dyevoich, N.S. Disher, M.A. Haro, and K.M. Haas, A TLR4-TRIF-dependent signaling pathway is required for protective natural tumor-reactive IgM production by B1 cells. Cancer Immunol Immunother 69 (2020) 2113–2124.

[22] F. Costa, J.E. Hagan, J. Calcagno, M. Kane, P. Torgerson, M.S. Martinez-Silveira, C. Stein, B. Abela-Ridder, and A.I. Ko, Global Morbidity and Mortality of Leptospirosis: A Systematic Review. PLoS Negl Trop Dis 9 (2015) e0003898.

[23] Y. Nagai, S. Akashi, M. Nagafuku, M. Ogata, Y. Iwakura, S. Akira, T. Kitamura, A. Kosugi, M. Kimoto, and K. Miyake, Essential role of MD-2 in LPS responsiveness and TLR4 distribution. Nat Immunol 3 (2002) 667–72.

[24] S.N. Vogel, C.T. Hansen, and D.L. Rosenstreich, Characterization of a congenitally LPS-resistant, athymic mouse strain. J Immunol 122 (1979) 619–22.

[25] A. Poltorak, X. He, I. Smirnova, M.Y. Liu, C. Van Huffel, X. Du, D. Birdwell, E. Alejos, M. Silva, C. Galanos, M. Freudenberg, P. Ricciardi-Castagnoli, B. Layton, and B. Beutler, Defective LPS signaling in C3H/HeJ and C57BL/10ScCr mice: mutations in Tlr4 gene. Science 282 (1998) 2085–8.

[26] N. Nair, and M. Gomes-Solecki, A Mouse Model of Sublethal Leptospirosis: Protocols for Infection with Leptospira Through Natural Transmission Routes, for Monitoring Clinical and Molecular Scores of Disease, and for Evaluation of the Host Immune Response. Curr Protoc Microbiol 59 (2020) e127.

